# Functional Electrical Stimulation of the Soleus Redistributes Lower-Limb Joint Work Distally in Young and Older Adults

**DOI:** 10.1101/2025.10.02.680079

**Authors:** Ningzhen Zhao, Ross H. Miller, Lisa Griffin, Owen N. Beck

## Abstract

Older adults walk with reduced ankle and greater hip mechanical output compared to young adults. This “distal-to-proximal redistribution” likely contributes to the greater metabolic energy expenditure during walking in older versus young adults. Due to the inverse relationship between ankle and hip use, functional electrical stimulation (FES) of the ankle extensors may increase ankle mechanical work and indirectly decrease hip mechanical work. Although FES increases stimulated muscle metabolism, bilateral soleus stimulation may restore more youthful walking kinetics without a detectable change in whole-body metabolism because ankle extension requires less metabolic energy than hip extension. Ten young adults and 10 older adults walked on a treadmill at 1.25 m/s with and without FES bilaterally applied over the respective leg’s soleus when the anterior-posterior ground reaction force exceeded 10% body weight. FES use altered walking mechanics and metabolic power similarly across age groups (all FES condition and age-group interactions p≥0.214). Across age groups, FES increased ankle mechanical power (p=0.041) and redistributed mechanical work production to occur relatively more at the ankle and less at the hip (p=0.010). The lower-limb joint redistribution ratio of older adults walking with FES was not different to that of young adults during baseline (p=0.785). Moreover, walking with FES increased metabolic power by 2% (p=0.037). FES attenuated older adult distal-to-proximal redistribution and modestly increased whole-body metabolic rate. FES applied to soleus muscles during the late stance of walking affects users similarly across the lifespan, indicating that FES interventions ought to consider a person’s functional needs, regardless of age.

## Introduction

During walking, older adult ankle kinetics have lower magnitudes than those of young adults. Older adults produce 9-12% lower peak ankle extensor moments (Anderson & Madigan, 2014; Winter et al., 1990) and 11–35% less peak ankle mechanical power (Beijersbergen et al., 2017; DeVita & Hortobagyi, 2000; Franz & Kram, 2013, 2014; Winter et al., 1990) compared to young adults at the same walking speed. Reduced ankle extensor kinetics are typically accompanied by an increase in hip extensor kinetics during walking (Delabastita et al., 2021; DeVita & Hortobagyi, 2000; Franz & Kram, 2013; Lewis & Ferris, 2008). This “distal-to-proximal redistribution” of joint kinetics in older adults likely contributes to their greater metabolic energy expenditure during walking compared to young adults (Delabastita et al., 2021). That is because hip extensors expend more metabolic energy to produce joint kinetics than ankle extensors (Fallah & Beck, 2025).

Neural strategies, not muscle capacity, may govern the diminutive ankle kinetics during walking in older adults. Generally, older adults retain the ability to produce youthful ankle kinetics during walking. For example, older adults can walk with more youthful ankle mechanical power and work magnitudes when instructed verbally or following visual feedback (Browne & Franz, 2018, 2019; Lewis & Ferris, 2008) and they naturally increase their ankle mechanical work >40% when walking uphill versus level ground (Franz & Kram, 2014). Even older adults with mobility impairments naturally increase their ankle mechanical power and work to walk at faster speeds (Graf et al., 2005; Kerrigan et al., 1998; Riley et al., 2001; Silder et al., 2008). While older adult ankle strength exhibits lower-to-moderate associations with ankle kinetics during walking (Delabastita et al., 2021; Hortobágyi et al., 2015; Judge et al., 1996; Silder et al., 2008), strength training typically does not restore more youthful preferred walking speeds or ankle kinetics in older adults (Beijersbergen et al., 2013, 2017; Symons et al., 2005). Perhaps directly intervening on the nervous system with an external stimulus can increase the mechanical output of older adult ankle extension during walking.

Older adults may be able to restore more youthful walking kinetics using wearable electrical stimulators. Supplementing voluntary ankle extension during walking with functional electrical stimulation (FES) can increase ankle mechanical power and work (Aout et al., 2024, Aout et al., 2025) by recruiting additional motor units (Bickel et al., 2011; Knaflitz et al., 1990). While FES during walking is rarely used by neurotypical adults due to the prospect of increasing metabolic energy expenditure (Chen et al., 2022; Muthalib et al., 2016; Watanabe et al., 2019; Ye et al., 2024a), the strategic use of FES applied to ankle extensors during walking may not substantially increase whole-body metabolic energy expenditure. Increasing late-stance ankle mechanical work, such as with FES applied to ankle extensors, likely reduces the subsequent limb’s collision losses (Huang et al., 2015) and decreases hip mechanical work (Aout et al., 2025; Browne & Franz, 2019; Fickey et al., 2018; Huang et al., 2015). Because hip extensors expend more metabolic energy to produce joint kinetics than ankle extensors (Fallah & Beck, 2025), the metabolic increase of FES applied to ankle extensors may be blunted by the respective decrease of hip extensor kinetics. Altogether, electrically stimulating ankle extensors of older adults may restore more youthful walking kinetics, without substantially increasing whole-body metabolism.

The primary goal of this study was to determine the mechanical and metabolic effects of stimulating older adult soleus muscles during the late stance of walking. To accomplish this goal, we compared walking mechanics, hip extensor muscle activation, and whole-body metabolic energy expenditure of older adults with and without FES applied to bilateral soleus muscles during late stance. We hypothesized that electrically stimulating soleus muscles during walking would shift lower-limb mechanical work production to occur relatively more at the ankle and less at the hip in older adults, without detectably changing whole-body metabolic energy expenditure compared to walking without stimulation. Additionally, we sought to determine whether advanced age influenced the mechanical and metabolic effects of walking with FES by testing young adults in addition to older adults.

## Methods

### Participants

Ten young adults (5 females; Avg ± SD; age: 21.2 ± 2.0 years; height: 1.75 ± 0.09 m; mass: 77.9 ± 14.2 kg) and ten older adults (5 females; age: 77.4 ± 5.5 years, height: 1.70 ± 0.10 m, mass: 73.3 ± 15.4 kg) participated in this study. Participants self-reported as being free of metabolic and cardiovascular disease, neurologic pathology, orthopedic disability, and not having a cardiac pacemaker or electronic implant. Individuals provided informed written consent prior to participation in accordance with the University of Texas at Austin’s Institutional Review Board.

### Protocol

Participants were asked to arrive at the lab at least three hours after last consuming calories. Upon arrival, we placed reflective markers and surface electromyography (EMG) sensors to the skin (or clothing for some reflective markers) of participants using double-sided tape. Specifically, we placed reflective markers to the skin and clothing of participants in accordance with the full-body Plug-in-Gait model and superficial to the following landmarks: (bilateral) anterior superior iliac spine, posterior superior iliac spine, mid-thigh, lateral knee-joint center, mid-shank, lateral ankle-joint center, posterior heel, base of the 2nd metatarsal head, lateral anterior head, lateral posterior head, acromioclavicular notch, mid-upper arm, elbow-joint center, mid-forearm, medial and lateral wrist-joint center, head of 2nd metacarpals; (single site) 7th cervical vertebra, 10th thoracic vertebra, clavicle, sternum, and right inferior angle of the scapula (Davis et al., 1991).

Next, we shaved and prepared the skin superficial to the participant’s gluteus maximus on both legs following SENIAM recommendations prior to EMG sensor placement (Hermens et al., 2000). We recorded gluteus maximus EMG signals during a maximal voluntary contraction as the participant stood and performed isometric hip extension (2000 Hz; Trigno, Delsys Inc., Boston, MA, USA).

Participants performed four, five-minute treadmill walking trials at 1.25 m/s on an instrumented split-belt treadmill (Motek Medical BV, Amsterdam, NL). The initial walking trial served as familiarization to treadmill walking. The second trial was used for the baseline walking assessment without FES. The third trial served as familiarization for walking with bilateral soleus electrical stimulation and to determine the stimulation intensity for the fourth trial. The fourth trial was performed with FES applied to the soleus muscles during late stance. A five-minute seated rest preceded each treadmill trial.

Between the second and third treadmill walking trials, we outfitted each participant with self-adhesive stimulation electrodes. We placed the anode over the soleus muscle, distal to the gastrocnemius, and the cathode 2 cm distal to the anode. Each stimulation electrode was 5.08 cm^2^ and tethered to a DS7A stimulator (Digitimer, Cambridge, UK). We stimulated the skin superficial to the uni-articular soleus and not the bi-articular gastrocnemius because we wanted to promote ankle extension without additional knee flexion. During the third and fourth walking trials, we stimulated the participant’s leg when the respective limb produced an anterior-posterior ground reaction force that exceeded 10% of the participant’s body weight. A custom Spike 2 (Cambridge Electronic Design, Cambridge, UK) script was used to trigger stimulation based on real-time ground reaction force data (Fig. 1). All stimulation trains were 30 Hz with pulse widths of 200 μs (Gregory et al., 2007; Ibitoye et al., 2016; Tan et al., 2014; You et al., 2014; Zheng et al., 2018). Prior to the third walking trial, the stimulation intensity was set to the participant’s highest tolerable stimulation intensity. During the third walking trial, we asked participants if the stimulation intensity could be increased at the first and third minutes and tuned the stimulation intensity accordingly. The stimulation intensity achieved at minute three of the third trial was used throughout the entire fourth walking trial and to quantify the effects of FES during walking.

**Figure 1.**
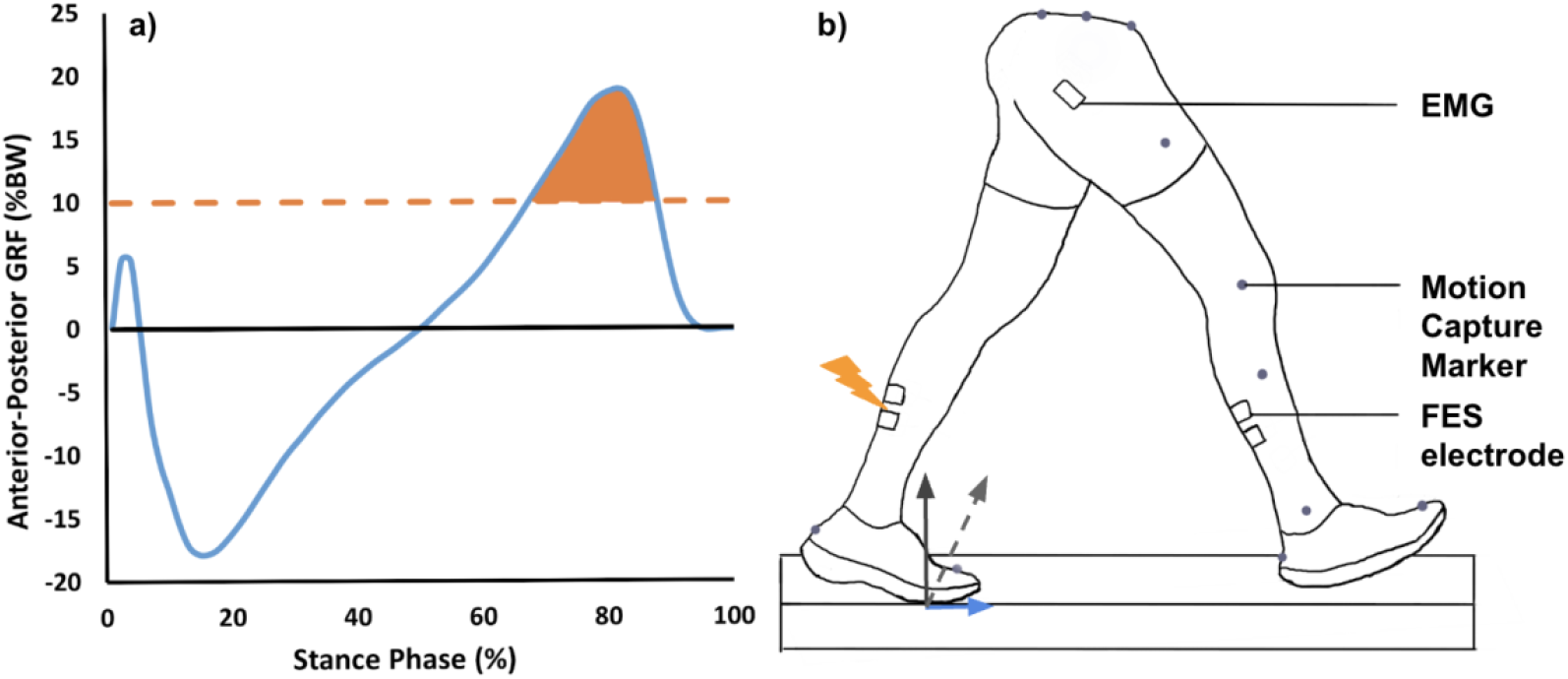
Participants received soleus functional electrical stimulation (FES) when the respective limb’s anterior-posterior ground reaction force (GRF) was >10% of their body weight (BW). a) Anterior-posterior GRF as a percentage of BW versus the stance-phase of walking. The orange dashed line represents the stimulation threshold (10% BW). b) Participant walking on a dual belt force-measuring treadmill, with a gluteus maximus electromyography (EMG) sensor, motion capture markers, and FES electrodes superficial to bilateral soleus muscles.

### Data collection and analyses

We recorded ground reaction forces, marker positions, and EMG signals during at least 12 steps over the last 30 seconds of each walking trial (Vicon Motion Systems, Oxford, UK). We averaged all steps within each condition for each participant. Ground reaction forces (2,000 Hz) were filtered using a 4th-order low-pass Butterworth filter (20 Hz). Marker trajectories were filtered using a Butterworth low-pass filter with a cutoff frequency of 6 Hz. We used filtered data to compute sagittal plane ankle and hip joint angles, angular velocities, moments, mechanical power, and mechanical work using Visual 3D (C-Motion, Inc., Germantown, MD, USA). We identified ground contact when the filtered vertical ground reaction force was >30 N. Subsequently, we calculated stride kinematic variables (duty factor, step and stride time), peak vertical and horizontal ground reaction forces, and select stance-phase joint mechanics (peak ankle moment, ankle power, positive ankle work, hip moment, hip power, positive hip work) using a custom MATLAB script (Mathworks, Natick, MA, USA).

To objectively quantify participant ankle versus hip mechanical use, we used participant redistribution ratio (Eq. 1) (Browne & Franz, 2018).

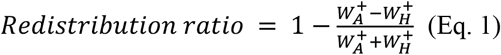

W_A_^+^ represents the positive mechanical work performed by the ankle per step, and W_H_^+^ represents the positive mechanical work performed by the hip per step. The redistribution ratio ranges from 0 to 2; 0 indicates that positive mechanical work was performed at the ankle but not the hip, and 2 indicates that mechanical work was performed at the hip but not ankle (Browne & Franz, 2018).

During each five-minute trial, we recorded the gas expired by participants using open-circuit spirometry (Parvo Medics, Sandy, UT). We averaged the rates of oxygen consumption 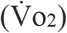 and carbon dioxide production 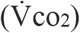 over the last 2 minutes of each trial and calculated mass-specific metabolic power (W/kg) using a standard equation (Péronnet & Massicotte, 1991) and dividing by body mass (W/kg).

We band-pass filtered gluteus maximus EMG signals at 20–500 Hz using a 2^nd^-order Butterworth filter and used a 60 Hz notch filter. We studied the largest hip extensor muscle’s EMG signals (gluteus maximus) due to its influence on participant hip extension. After rectifying the signals, we applied a 40 ms moving average window. We scaled each muscle’s EMG signal during walking to the peak EMG signal amplitude from the MVC trial. The peak EMG amplitude was defined as the highest 500 ms average within the MVC trial. We calculated peak, average, and integrated EMG signals for the gluteus maximus during walking using a custom MATLAB script.

### Statistical Analysis

We conducted all statistical analyses using SPSS (v28.0, IBM) with a significance threshold of α=0.05. We performed two-way repeated-measures ANOVAs with independent variables (age group and FES condition) versus neuromechanical and dependent variables (peak vertical ground reaction force, peak anterior posterior ground reaction force, duty factor, step time, peak ankle moment, peak ankle power, positive ankle work, peak hip moment, peak hip power, positive hip work, redistribution ratio, average gluteus maximus EMG, peak gluteus maximus EMG, integrated gluteus maximus EMG, and gross metabolic power). If there was no interaction between age group and FES use on the respective dependent variable, we performed two one-way ANOVAs to evaluate independent relationships between age groups and FES use on the respective dependent variable. If there was a significant interaction between our independent variables, we conducted post hoc comparisons to identify differences between conditions within each age group and between age groups within each condition. To mitigate the chance of reporting a Type I error across the pairwise comparisons, we applied a Bonferroni correction, setting the significance threshold at α=0.0125 (0.05/4). Because we were interested in the possibility of restoring youthful walking redistribution ratios in older adults with FES use, we also performed a t-test that compared the baseline Redistribution Ratio in young adults vs. the redistribution ratio during FES in older adults.

## Results

FES applied to the soleus during late stance reduced the redistribution ratio during walking, indicating that participants performed relatively more ankle work and less hip work compared to baseline. There were no interactions between age group and FES condition on any dependent variable pertaining to joint mechanics (all p≥0.214). Therefore, we interpreted the influence of FES condition on walking mechanics across age groups unless otherwise stated. Peak ankle moments and mechanical power were 4-6% greater during walking with FES versus baseline (p=0.050 and p=0.041, respectively) (Fig. 2). Peak hip moments and mechanical power were numerically 3-6% lower in the FES versus baseline condition, despite not achieving statistical significance (both p≥0.105) (Fig. 2). FES increased positive ankle work 8% compared to baseline (p=0.007), and decreased positive hip work 13% (p=0.070). Overall, these FES-induced joint-level changes decreased redistribution ratio 13% versus baseline (p=0.010) (Fig. 3). As a result, older adult redistribution ratio during walking with FES was indistinguishable from that of young adult baseline (p=0.785) (Fig 3).

**Figure 2.**
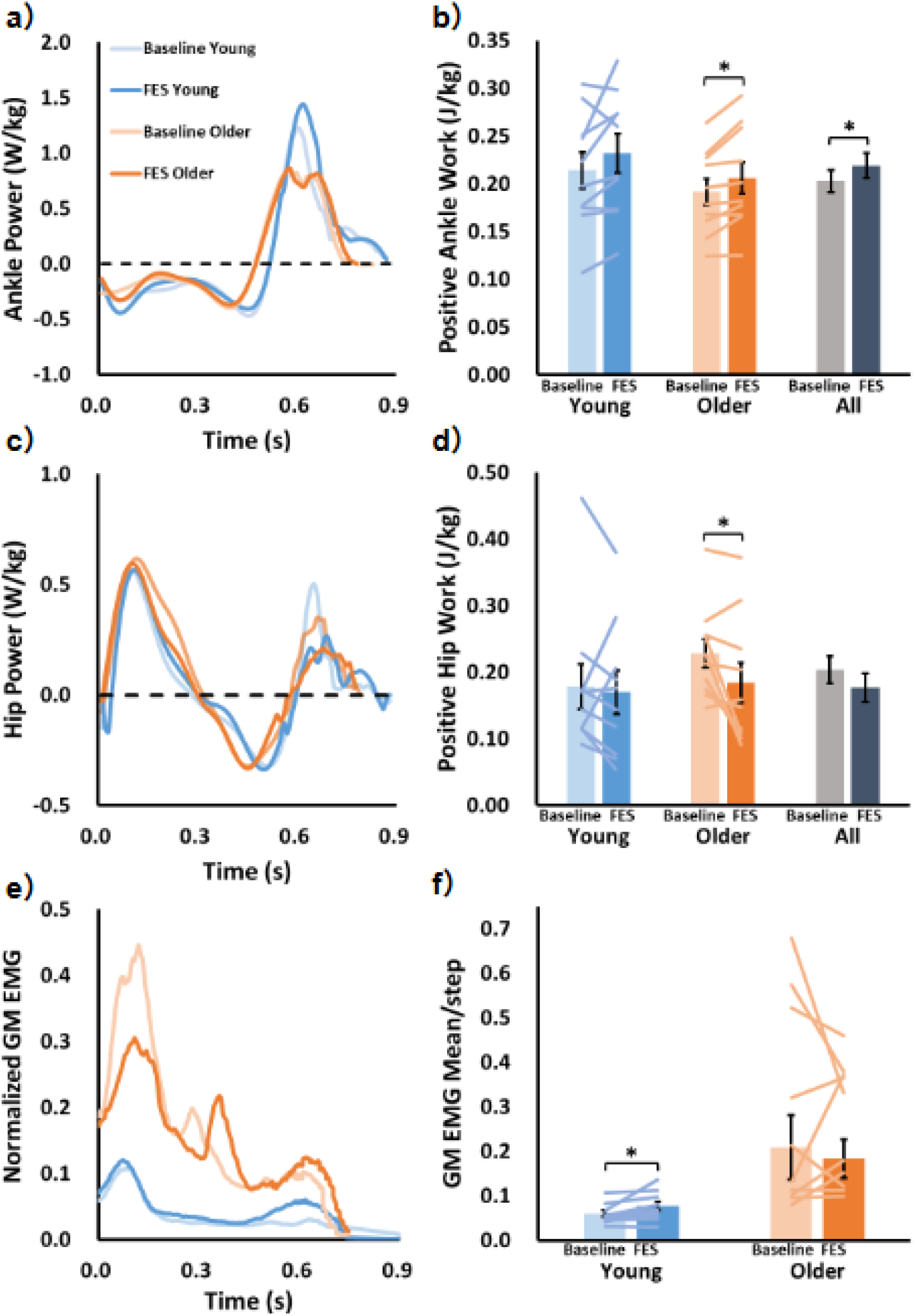
Joints kinetics and gluteus maximus EMG signal at baseline and FES conditions. Averages for each experimental condition are shown in the left column. Bar graphs in the right column represent group average (±SE) with individual data represented for each participant. a) Ankle power, c) hip power, and e) scaled gluteus maximus (GM) EMG signals for both age groups and FES conditions. b) Positive ankle work, d) Positive hip work, and f) average GM EMG per step in young and older adults with and without FES as well as across age groups. Data represent young adults (blue), older adults (orange), and all participants (gray). Lighter shades show baseline; darker shades show FES. Asterisks (*) indicates that p value < 0.05.

**Figure 3.**
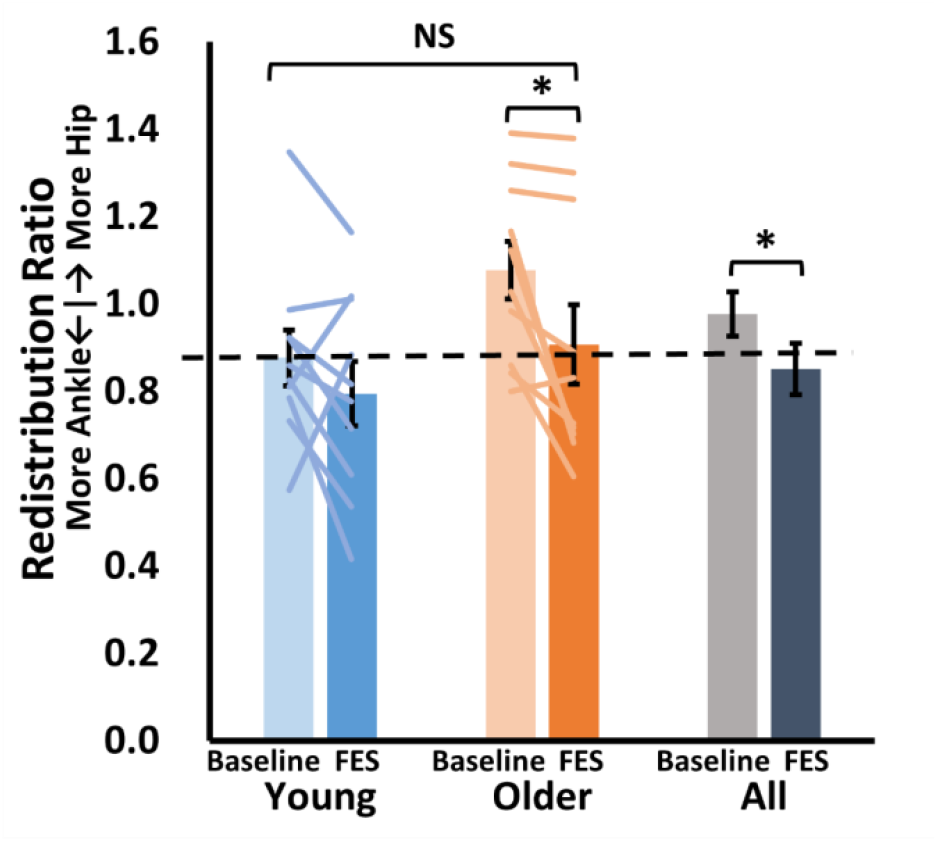
Functional electrical stimulation (FES) reduced redistribution ratio versus baseline. Average (±SE) redistribution ratio for young adults (blue), older adults (orange), and all participants (gray). Lighter shades show baseline; darker shades show FES. Dashed line represents the average baseline redistribution ratio of young adults. Asterisks (*) indicates that p value < 0.05; NS indicates that p value ≥ 0.05.

FES use altered participant ground reaction force profile without detectable changes in kinematics (Table 1). Notably, FES increased peak anterior-posterior GRF by 3% (p=0.036) and there were no detectable changes in step time or duty factor (both p≥0.475) (Table 1).

**Table 1.**
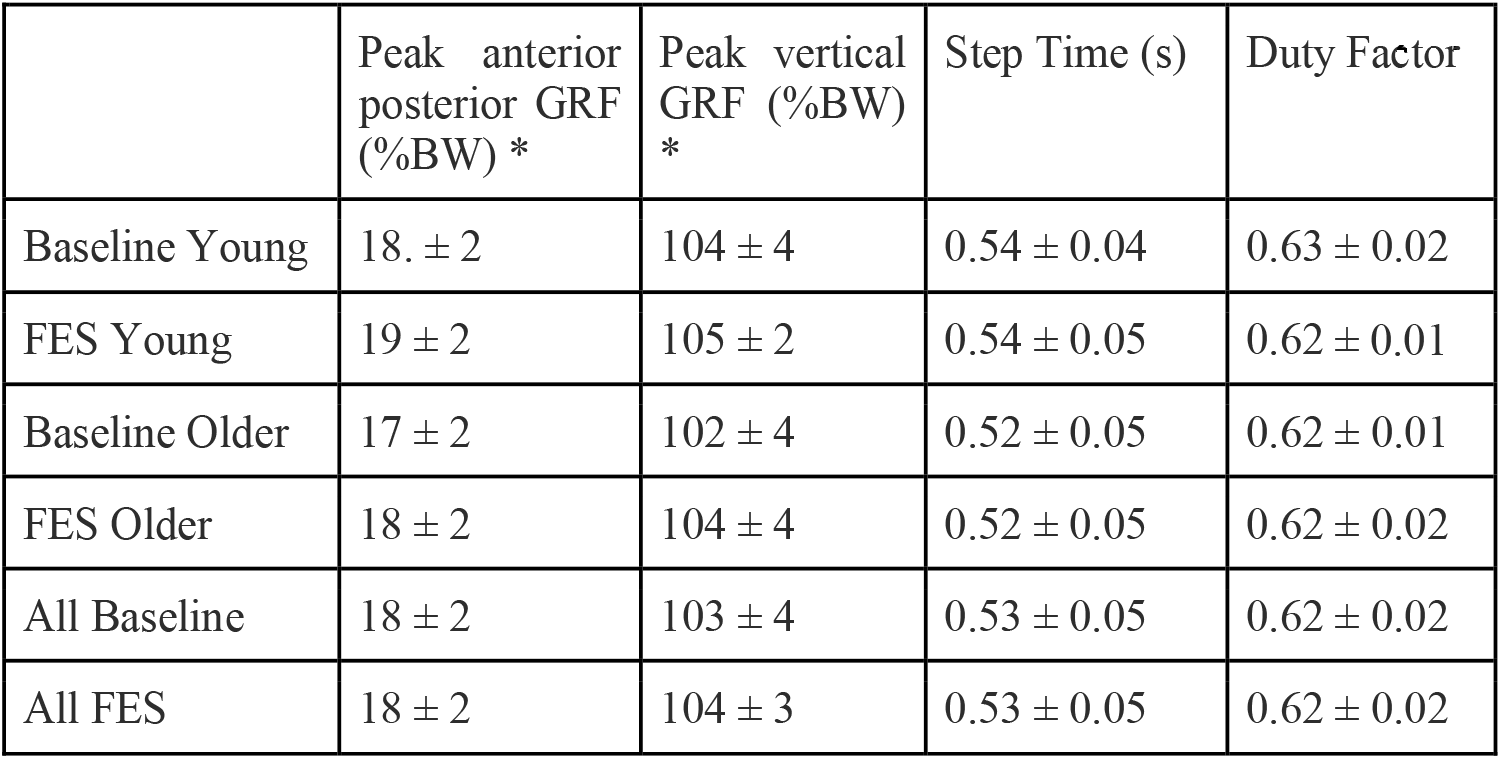
Average ± SD values of ground reaction force (GRF) metrics and kinematics during baseline and FES conditions, with significant main effects indicated by asterisks (*) (p<0.05).

Consistent with the established age-related increase in metabolic rate (Malatesta et al., 2002; Mian et al., 2006; Hortobágyi et al., 2011), our older participants expended 15% more gross metabolic power during walking than young adults at baseline (p=0.017). There was no interaction between FES use and age group on gross metabolic power during walking (p=0.527). Walking with FES increased participant metabolic power 2% versus baseline across age groups (p=0.037) (Fig. 4).

**Figure 4.**
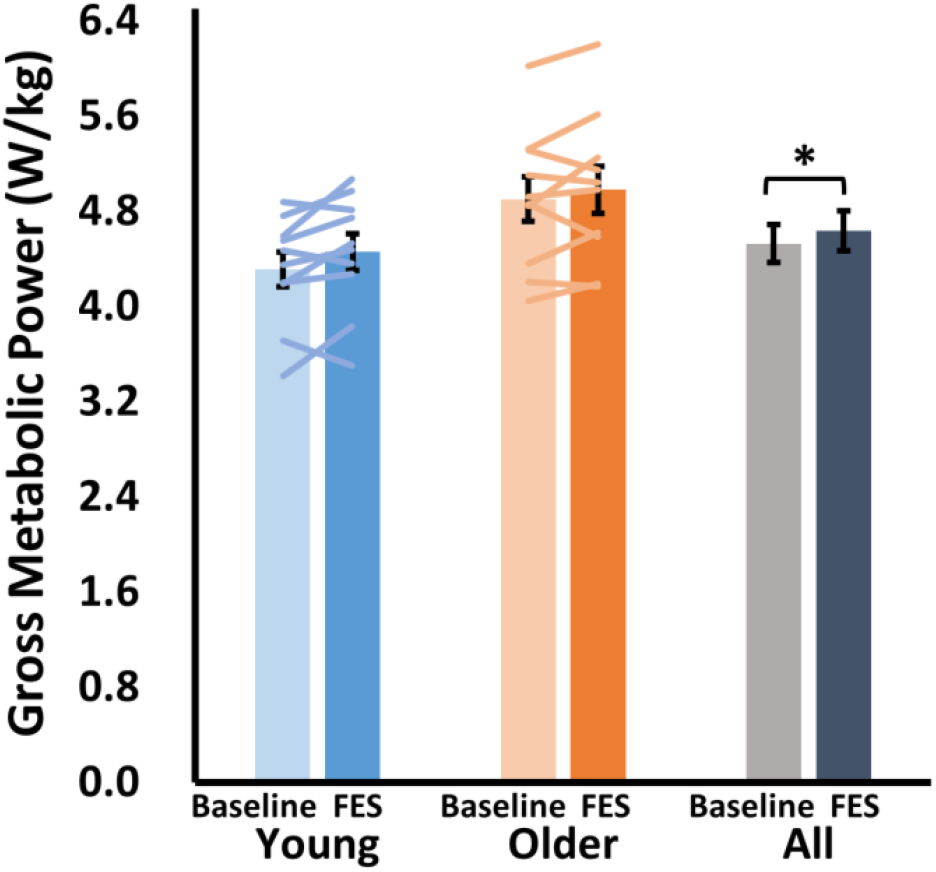
Walking with functional electrical stimulation (FES) increased participant metabolic power versus baseline. Average (±SE) net metabolic power for young adults (blue), older adults (orange), and all participants (gray). Lighter shades show baseline; darker shades show FES. Asterisks (*) indicates that p value < 0.05.

Age group and FES use did not interact, nor did either independent variable relate to peak, average or integrated gluteus maximus EMG signals (all p≥0.369) (Fig. 2).

## Discussion

To mitigate the age-related distal-to-proximal redistribution in walking kinetics (DeVita & Hortobagyi, 2000), we applied FES to participant ankle extensors during late stance. By applying FES over a functionally relevant and brief period, we increased ankle mechanical output to the extent that older adults with FES exhibited a non-different redistribution ratio compared to young adult baseline. Concurrently, whole-body metabolic energy expenditure increased 2% during walking with FES versus baseline, rejecting our hypothesis. Thus, the metabolic increase due to muscle stimulation (Chen et al., 2022; Muthalib et al., 2016; Watanabe et al., 2019; Ye et al., 2024b) trumped the potential decrease due to the altered joint kinetics (Delabastita et al., 2021; Fallah & Beck, 2025). Moreover, late-stance soleus FES affected the walking mechanics and metabolism of neurotypical young and older adults similarly. Therefore, researchers and healthcare professions may not need to consider to the physiological changes associated with advanced age when devising acute FES-walking interventions.

The effects of applying FES to older adult leg muscles during walking support contemporary views regarding the age-related distal-to-proximal redistribution. Older adults characteristically walk using their ankles relatively less and hips relatively more than young adults (DeVita & Hortobagyi, 2000; Boyer et al., 2017; Delabastita et al., 2021). By using FES to increase participant ankle mechanical work, our results corroborate the notion that older adults can walk with more youthful ankle kinetics (Browne & Franz, 2018, 2019; Franz, 2016; Lewis & Ferris, 2008). Our results also coincide with evidence that ankle mechanical work inversely relates with hip mechanical work during walking (Browne & Franz, 2018, 2019; Delabastita et al., 2021). As such, interventions that target the older adult nervous system, including the use biofeedback to increase neural drive (Browne & Franz, 2018, 2019; Lewis & Ferris, 2008), may be more effective at mitigating the age-related distal-to-proximal redistribution in joint kinetics than those that primarily target older adult muscle-tendon mechanics.

We were unable to control for a potential trial order effect. Due to potential short-term carryover effects of FES during walking (Kido Thompson & Stein, 2004; Palmer et al., 2017; Thompson et al., 2006), we did not reassess baseline walking after the FES trials. Hence, we were unable to control for the potential influence of trial order on our results. Notably, we conducted familiarization trials prior to the baseline and FES experimental trials, and we provided at least 5 minute of rest between walking trials to mitigate the potential effects of fatigue. Still, it is possible that fatigue increased throughout our protocol, which is common with the use of electrical stimulation (Boerio et al., 2005; Jones et al., 1979; McNeil et al., 2006). However, the potential effect of fatigue does not affect our conclusions. That is because greater fatigue in later trials would have likely mitigated the effects of FES on ankle joint kinetics and further increased participant metabolism compared to baseline (Allen et al., 2008; Pethick & Tallent, 2022; Sahlin, 1992). In other words, if our participants were more fatigued during later trials, the ability of FES use to acutely restore more youthful walking kinetics and metabolism in older adults may have been underreported.

Withstanding the physiological changes that occur over 57 years (Boss & Seegmiller, 1981; Preston & Biddell, 2021), FES modified joint kinetics and metabolic energy expenditure similarly for young and older adults. While potential functional outcomes and long-term physiological adaptations may differ across age groups, our acute intervention indicates that FES paradigms should be prescribed based on a person’s biomechanical needs, regardless of age.

## Ethics approval and consent to participate

Participants provided informed written consent as per the University of Texas Institutional Review Board’s approved protocol (STUDY00004713).

## Consent for publication

Not applicable.

## Availability of data and materials

The dataset supporting the conclusions of this article is included within the article.

## Authors’ contributions

N.Z. contributed to study conception, data acquisition, interpretation of data, manuscript draft, and approved of the final draft. R.H.M. contributed to study conception, interpretation of data, providing manuscript feedback and final draft approval. L.G. contributed to study conception, interpretation of data, manuscript editing, and approved of the final draft. O.N.B. contributed to study conception, data acquisition, interpretation of data, manuscript draft and editing, and approved of the final draft. All authors agree to be personally accountable for the author’s own contributions and to ensure that questions related to the accuracy or integrity of any part of the work.

## Acknowledgements

Not applicable.

